# CHRRT: boosting coordinate hit-and-run with rounding by thinning

**DOI:** 10.1101/2022.11.17.516802

**Authors:** Johann F. Jadebeck, Wolfgang Wiechert, Katharina Nöh

## Abstract

Thinning is a sub-sampling technique to reduce the memory footprint of Markov chain Monte Carlo. Despite being commonly used, thinning is rarely considered efficient. For sampling convex polytopes uniformly, a highly relevant use-case in systems biology, we here demonstrate that thinning generally boosts computational and, thereby, sampling efficiencies of the widely used Coordinate Hit-and-Run with Rounding (CHRR) algorithm. We benchmark CHRR with thinning (CHRRT) with simplices and constrained-based metabolic networks with up to thousands of dimensions. With appropriate thinning, CHRRT offers a substantial increase in computational efficiency compared to unthinned CHRR, in our examples of up to three orders of magnitude, as measured by the effective sample size per time (*ESS/t*). Our experiments reveal that the performance gain of CHRRT by optimal thinning grows substantially with polytope (effective model) dimension. Based on our experiments, we provide practically useful advice for tuning thinning to efficient and effective use of compute resources. Besides allocating computational resources optimally to permit sampling convex polytopes uniformly to convergence in a fraction of time, exploiting thinning unlocks investigating hitherto intractable models under limited computational budgets. CHRRT thereby paves the way to keep pace with progressing model sizes within the existing constraint-based reconstruction and analysis (COBRA) tool set. Sampling and evaluation pipelines are available at https://jugit.fz-juelich.de/IBG-1/ModSim/fluxomics/chrrt.

## 1 Introduction

Convex polytopes are geometrical objects that constrain the parameter space in many applied modeling contexts, including operations research (Ciomek and Kadziński, 2021), ecological modeling (Drouineau et al., 2021), computational finance (Chalkis et al., 2021a), astronomy (Lubini and Coles, 2012), physics (Leake et al., 2020), and systems biology (Herrmann et al., 2019). Uniform convex polytope sampling (UCPS), i.e., drawing representative random numbers from a truncated uniform distribution defined over a bounded polytope, has become a standard tool for model analysis, parameter space exploration/characterization, and volume approximation. UCPS has a variety of applications, especially in systems biology, where mass balances under steady state conditions together with physiological constraints induce convex polytopes (De Martino et al., 2015; Theorell and Stelling, 2022). Typical use cases include the unbiased characterization of the solution spaces of metabolic (Schellenberger and Palsson, 2009; Loghmani et al., 2022) and gene networks (Machado et al., 2016), identification of metabolic flux couplings (Heinonen et al., 2019), design of experiments (Schellenberger et al., 2012; Beyß et al., 2021), and assessing the effects of uncertainties in biochemical network formulations (Bernstein et al., 2021; Dinh et al., 2022). Here, genome-scale models (GEMs) are routinely reconstructed from genomic and biochemical information (Robinson et al., 2020), resulting in comprehensive model formulations with high-dimensional polytopes as solution spaces. For instance, as of now the largest metabolic model in the BiGG database (King et al., 2015; Norsigian et al., 2019), *Recon3d* (Brunk et al., 2018), accounts for more than ten thousand reactions and has a polytope dimension of 4, 861 (median dimension of BiGG models: 594). Current constraint-based reconstruction and analysis (COBRA) modeling initiatives, targeting the description of microbial communities or the human body (Colarusso et al., 2021; Thiele et al., 2020; Heinken et al., 2021), are escalating GEM sizes further.

A simple UCPS strategy is *rejection sampling*, a Monte Carlo technique that iteratively generates samples within easily accessible objects such as polytope-enclosing (hyper)cubes, where it keeps only those draws that are located interior to the polytope, until the truncated uniform target distribution is approximately covered (Gelman et al., 2013). While initial efforts in the field of systems biology used this technique (Wiback et al., 2004), rejection sampling was quickly recognized to be computationally infeasible for more comprehensive network models, because for every feasible sample the number of rejections grows super-exponentially with polytope (effective model) dimension.

To overcome this bottleneck, the class of Hit-and-Run (HR) algorithms has been employed (Smith, 1984). HR is a Markov chain Monte Carlo (MCMC) technique, which, starting from an initial flux state within the polytope, generates a sequence of flux states (a Markov chain) by drawing randomly from inner chords directed in random directions (Kaufman and Smith, 1998). As a result, the generated states, or a fraction of them, are returned as MCMC *samples*. These samples, albeit being correlated by construction, approximate the target distribution asymptotically (Lovász, 1999).

The critical metric for practical applicability of HR is the *sampling efficiency*, which measures how quickly the target distribution is approximated well in terms of wall-clock time. Sampling efficiency is determined by two components: (1) the computational efficiency of the HR sampling algorithm, i.e., how many computational operations are required per generated state, and (2) the statistical efficiency, which specifies how well a set of samples represents a target distribution. Formally, the statistical efficiency of *N* samples produced by a Markov chain is given by the *effective sample size* (*ESS*) (Gelman et al., 2013):

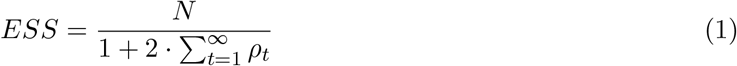

with *ρ*_*t*_ being the autocorrelation of the sequence at lag *t*. The sampling efficiency of the MCMC implementation is then measured by the *ESS* normalized to wall clock time *t*, i.e., *ESS/t*.

According to Eq. (1), two strategies to raise MCMC sampling efficiencies are (1) reducing the computational cost per sample, and (2) decreasing the autocorrelation between subsequent states. For HR, the cost per sample is reduced by updating the states in a pure coordinate-wise fashion instead of drawing random directions, a technique also known as Gibbs sampling (Robert and Casella, 2004). To decrease autocorrelation, the average distance between subsequent states is to be increased. In UCPS, the achievable distance is limited by the geometry of the polytope. Intuitively, in isotropic geometries on average longer steps are possible than in non-isotropic spaces. Since heterogeneous parameter scales in GEMs are the rule, an affine transformation is employed to homogenize the scales prior to sampling, the so-called *rounding transformation* (De Martino et al., 2015). After sampling, the states have to be mapped back to the original space by inverting the rounding transformation. Together, the two strategies culminated in the celebrated Coordinate Hit-and-Run with Rounding (CHRR) algorithm (Haraldsdóttir et al., 2017), the tried-and-tested workhorse for UCPS in the COBRA domain (Herrmann et al., 2019; Fallahi et al., 2020; Theorell et al., 2021), with several implementations being available (Haraldsdóttir et al., 2017; Heirendt et al., 2019; Jadebeck et al., 2020; Gollub et al., 2021).

In practice, a common MCMC post-processing technique is *thinning*, or sub-sampling, which refers to discarding all but a representative subset of the generated samples. The primary motivation for thinning is the reduction of the sampling memory footprint, which becomes crucial when investigating large-scale GEMs. For sampling *Recon3d* to convergence, as an example, CHRR requires *O* (10^9^) iterations, generating ∼80 TB of uncompressed data, which is clearly impractical for routine application. The most widely used form of thinning is *fixed frequency* thinning with thinning constant *τ*, which means that only every *τ*^th^ sample is kept. In constraint-based metabolic modeling, some studies report thinning constants of 100, 1000, and 10,000 (Herrmann et al., 2019; Fallahi et al., 2020), without providing guidance for the suitable selection of *τ* for the sampling problems at hand. On the other hand, Haraldsdóttir et al. (2017) pointed out that *τ* should be selected depending on the problem dimension. Specifically, the authors suggest setting *τ*, as a rule of thumb, to eight times the squared polytope dimension, stating that this choice ensures statistical independence of the thinned samples. Since CHRR produces highly correlated subsequent states, thinning indeed reduces the autocorrelation of the samples. Without thinning, however, CHRR produces a better approximation to the target distribution in the same number of iterations. With other words, thinning discards information, rendering the given argument statistically questionable. Indeed, thinning is known to be statistically inefficient in all but very few cases (Geyer, 1992; Link and Eaton, 2012).

In this work, we study the impact of thinning for CHRR systematically from the perspective of sampling efficiency, based on principled statistical criteria. By analyzing simplices and GEMs with widely varying dimensionality, we demonstrate that CHRR is one of the rare cases for which thinning generally boosts sampling efficiencies, measured in terms of *ESS/t*. From our benchmarks, we derive simple, yet effective and statistically rigorous rules to optimally tune CHRR thinning, which we verify using *Recon3D*. Our findings guide practical sampling efforts of COBRA models towards achieving more effective samples with less compute resources.

## 2 Methods

We start by giving relevant background about UCPS, describe established sampling procedures, test problems, and MCMC along with implementation details.

### 2.1 Pre- and post-processing convex GEM flux polytopes

GEMs are stoichiometric network models that are compiled from genomic information and biochemical knowledge (Kim et al., 2017). By mass balancing at steady-state, linear inequality systems are derivedfrom these models for the unknown metabolic reaction rates (fluxes) *ν*:

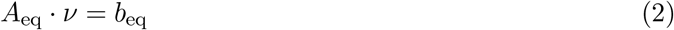

where *A*_eq_ is the (extended) stoichiometric matrix, and *b*_eq_ contains the time derivatives of the (intra- and extracellular) metabolite concentrations. In addition, fluxes are subject to linear inequalities originating from lower and upper limits on their values:

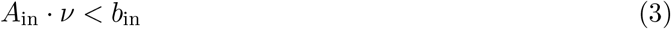

with *A*_in_ the constraint matrix and *b*_in_ contains the flux bounds for each flux. Together, Eqs. 2 and 3 define the *D*-dimensional flux polytope of a given GEM:

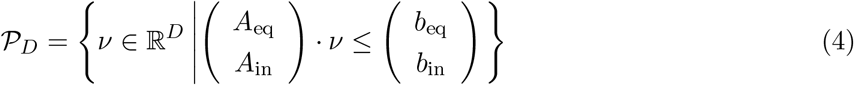

In a first pre-processing step, redundant inequalities are eliminated from Eq. (4). Then, inequalities that are effectively equalities are identified and reformulated as equalities. These manipulations leave the solution space of the flux polytope 𝒫_*D*_ unchanged, but improve numerical stability of the sampling workflow (Theorell et al., 2021). Next, the null space *K* of the (adapted) equality system is used to express the polytope in terms of orthogonal (independent) coordinates *ν*^indep^:

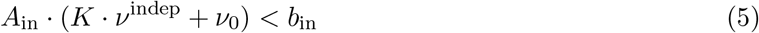

where *ν*_0_ is some flux solution within the flux polytope 𝒫_*D*_, and *ν*^indep^ ∈ ℝ^*d*^ with *d* ≤ *D* the *effective* model dimension. A typical choice for *ν*_0_ is the Chebyshev center of the flux polytope. By applying Eq. (5), all equalities are eliminated from 𝒫_*D*_ in Eq. (4), resulting in the dimension-reduced flux polytope 𝒫_*d*_:

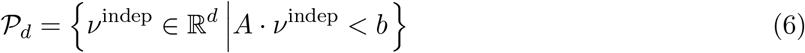

where *A* = *A*_in_ · *K* ∈ ℝ^*m* × *d*^ and *b* = *b*_in_ − *A*_in_ · *ν*_0_ ∈ ℝ^*m*^ with *ν*_0_ and *m > d*. As an example, *Recon3D*, as downloaded from BiGG, has 10, 600 bounded reactions. After dimension-reduction and flux coordinate transformation, *m* = 11, 195 inequality constraints and *d* = 4, 861 effective dimensions remain.

Next, the dimension-preserving linear rounding transformation *R* is determined, mapping a vector of independent fluxes *ν*^indep^ ∈ 𝒫_*d*_ to a transformed flux vector *ν*^round^ in a more isotropic flux polytope 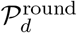. Specifically, after computing the Maximum Volume Ellipsoid (MVE) inscribed in 𝒫_*d*_ (Zhang and Gao, 2003), the inverse mapping *R*^−1^ is determined such that it maps the MVE to the unit sphere. The rounded polytope 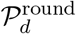 is then given by:

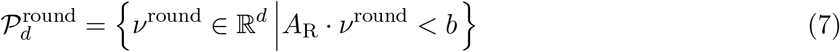

with *A*_R_ = *A* · *R*^−1^ ∈ ℝ^*m* × *d*^. Finally, any flux vector *ν*^round^ in the rounded space has to be mapped back to a flux vector *ν* in the original *D*-dimensional flux space by the affine back-transformation

*T* : *ν*^round^ 1 ↦ *ν* given by:

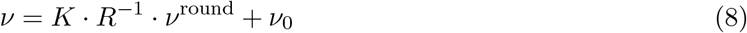

### 2.2 CHRRT algorithm

Thinning is an integral part of many HR algorithms. We hitherto denote implementations of CHRR that make deliberate use of thinning by *CHRRT*. Listing 1 gives a pseudocode description of the CHRRT algorithm. With a UCPS task at hand, defined by Eqs. (2)-(3), first the rounded polytope 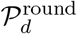is determined (L 1). Starting from an initial flux state, samples in 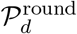 are iteratively generated by constructing chords along the coordinates (L 6-10). Fixed frequency thinning is applied with thinning constant *τ* (L 11). Lastly, every sample that is not discarded due to thinning is transformed back to the full dimensional flux space 𝒫_*D*_ (L 12).

#### Algorithm 1

**Coordinate Hit-and-Run with Rounding and Thinning (CHRRT)**

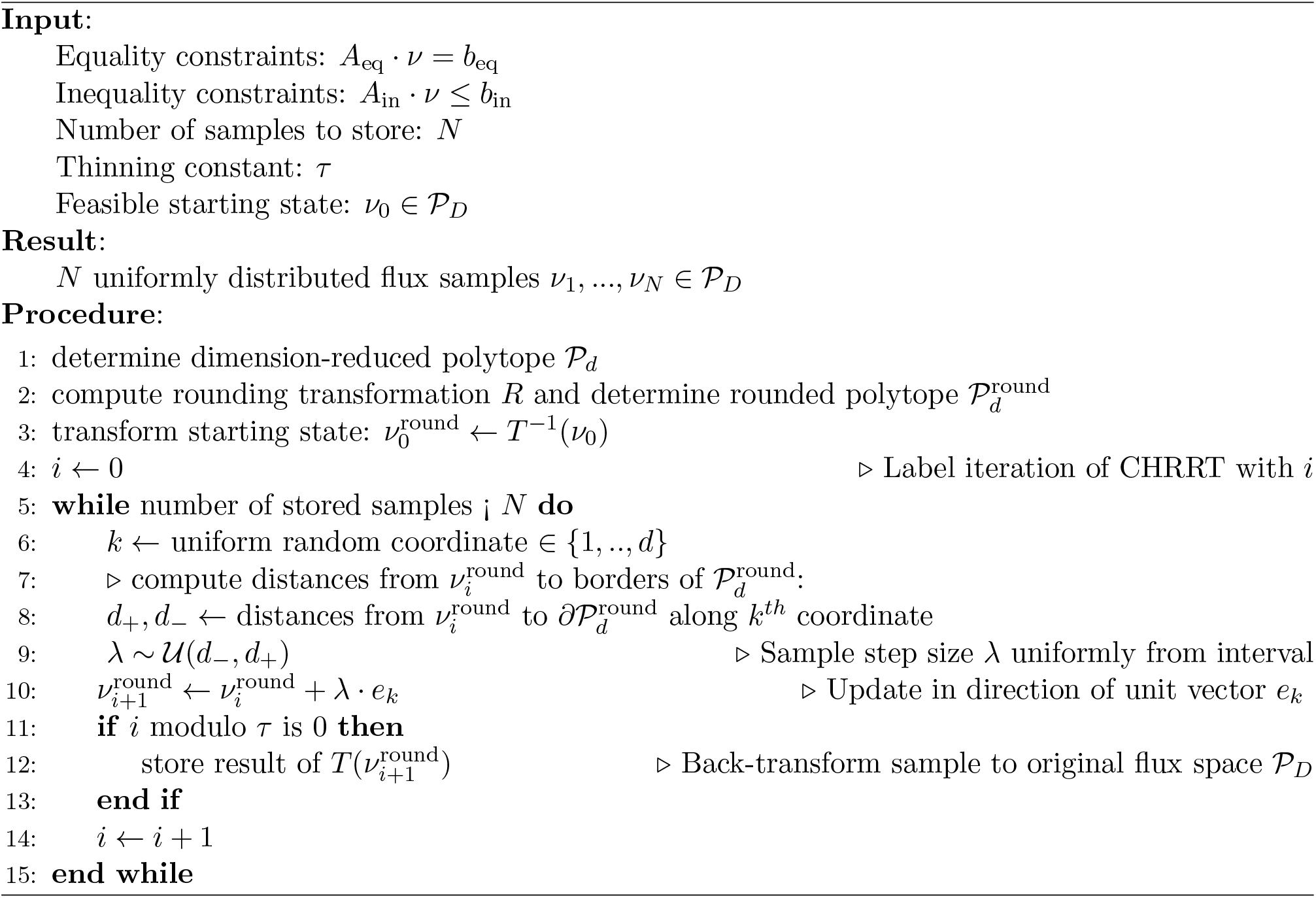

Concerning the computational costs of CHRRT, determining an internal point *ν*_0_ and computing the rounded polytope 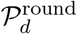 are one-time costs, both being independent of the numbers of samples. We therefore do not further consider these costs in this work. The most expensive operation per sample is the back-transformation *T* that maps the generated sample to the original polytope (Listing 1, line 12). The back-transformation amounts to a dense matrix-vector multiplication, with worst case costs of 𝒪(*m* · *d*) per sample. All other operations are element-wise additions, multiplications, or divisions with per-sample costs of 𝒪 (*d*).

### 2.3 Test problems

We consider two types of test problems, simplices representing well-defined and easy to scale problems, and GEMs being our main concern. A *d*-dimensional simplex 𝒮_*d*_ is defined by:

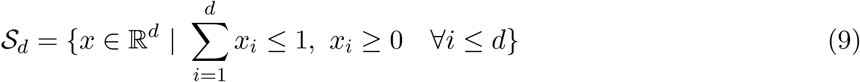

Being easily scalable to different dimensions, simplices are ideal to study the dependency of sampling complexity and polytope dimension. Different to GEMs, simplices are constructed by non-redundant constraints and, therefore, do not need to be dimension-reduced. However, as for GEMs, polytope rounding improves the geometric isotropy of simplices. Thus, Eq. (8) remains valid, with the matrix *K* set to identity. In this work, simplices in 64, 256, 1,024, and 2,048 dimensions were benchmarked. Besides simplices, we selected nine GEMs of varying sizes for this study, eight for benchmarking thinning, and one (*Recon3D*) for validation purposes. Effective polytope dimensions range from 24 to 4,861. In contrast to simplices, for GEMs there is no consistent relationship between the number of reactions, the number of constraints, and the effective polytope dimension *d*. We refer to SI Table S.1 for more details on the test problems.

### 2.4 MCMC convergence diagnostics

Assessing whether a (sub)sequence of MCMC samples has converged to the target distribution is not only vitally important in practical inference, but also critical for reliable benchmarking sampling performances. The *ESS* is computed for each of the *d* dimensions separately and set to the minimum value. Particular care must be taken here, since the estimated *ESS* is typically noisy and may therefore be unreliable for sample sets that are far from being converged. We here follow the advice given by Vehtari et al. (2021) to use four independent chains and compute the rank-normalized 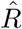 diagnostic along with the *ESS* to test for convergence. Vehtari et al. (2021) argue that the estimation of 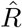 and *ESS* requires reliable estimates of the variances and autocorrelations of the samples, that is, the estimation of 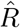 and *ESS* is only reliable, if the *ESS* of a set of samples is sufficiently large. Therefore, in our study, the threshold for convergence is set to 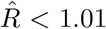 and *ESS >* 400, which is suitable to a setup with four independent Markov chains (the *ESS* scales linearly with the number of chains). Note that still each chain is required to converge.

### 2.5 Implementation details

GEMs were downloaded from the BiGG database (King et al., 2015; Norsigian et al., 2019) in SBML (Systems Biology Markup Language) format (Hucka et al., 2003), and plugged into PolyRound v0.1.8, a python package for polytope rounding (Theorell et al., 2021). The highly optimized polytope sampling library HOPS v3.0.0 was used for MCMC sampling (Jadebeck et al., 2020). CHRRT, as implemented in HOPS, was used for all benchmark runs to ensure their comparability. In each sampling run, four parallel chains were used. Samples, the rounded polytopes, and the accompanying transformations were stored. The 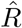 diagnostic and the *ESS* to test for MCMC convergence are calculated using arviz v0.12.1 (Kumar et al., 2019). All numerical experiments were run on an AMD Ryzen 9 5950X 16-Core Processor.

## 3 Results and discussion

After motivating the idea of using thinning in CHRRT to reallocate computational resources, we quantify achievable sampling efficiencies for the test problems. From that, we derive simple and useful calculation rules for near optimal thinning choice, which we compare to previous advice.

### 3.1 The role of thinning for CHRRT

Spurred by the seminal theoretic work of Geyer (1992), we set out to investigate whether the sampling efficiency of CHRRT benefits from thinning. The question is whether, for a fixed computational budget, CHRRT with *τ* > 1 convergences faster than unthinned CHRRT with *τ* = 1. To this end, we consider the *ESS* of a set of *N* samples in relation to the time cost of producing the samples, *t*_*N,τ*_. The time cost *t*_*N,τ*_ is given by the sum of the time costs *t*_update_ for producing *N* · *τ* states 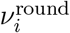 in Listing 1 (L 6-10), and the time cost *t*_transform_ of transforming the *N* samples back to the original flux space (Listing 1, L 12):

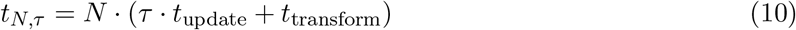

Geyer (1992) argued that for thinning to be beneficial, the time cost *t*_*N,τ*_ has to grow slower with *τ* than the *ESS*, indicating that thinning is not advantageous for every combination of sampling problem and MCMC algorithm. In this vein, Link and Eaton (2012) criticize the routine application of thinning for applications in ecology, where thinning is often detrimental to sampling efficiency.

To investigate the role of thinning for UCPS using CHRRT, we measured *t*_update_ and *t*_transform_ for a set of synthetic and real test problems (SI Table S.1). For both time costs, we took the wall-clock times and averaged them over the number of times each operation was applied. The resulting time ratios *t*_transform_*/t*_update_ are shown in Fig. 1. Among all test problems, the time ratios for only the two models with the fewest dimensions, i.e., the 64-dimensional simplex and the 24 dimensional *E. coli* core model, are below ten. For the remaining investigated models, *t*_transform_ is up to three orders of magnitude larger than *t*_update_. Precisely, for the simplices, the time ratio increases quadratically with dimension. We also observed a stark increase in the ratio for GEMs. Different to simplices, however, with GEMs additional factors contribute to the time ratios, such as the number of fluxes *D* (more fluxes increase *t*_transform_) and the number of constraints *m* (more constraints increase *t*_update_), resulting in a less succinct relation between effective model dimensions and time ratios.

**Figure 1:**
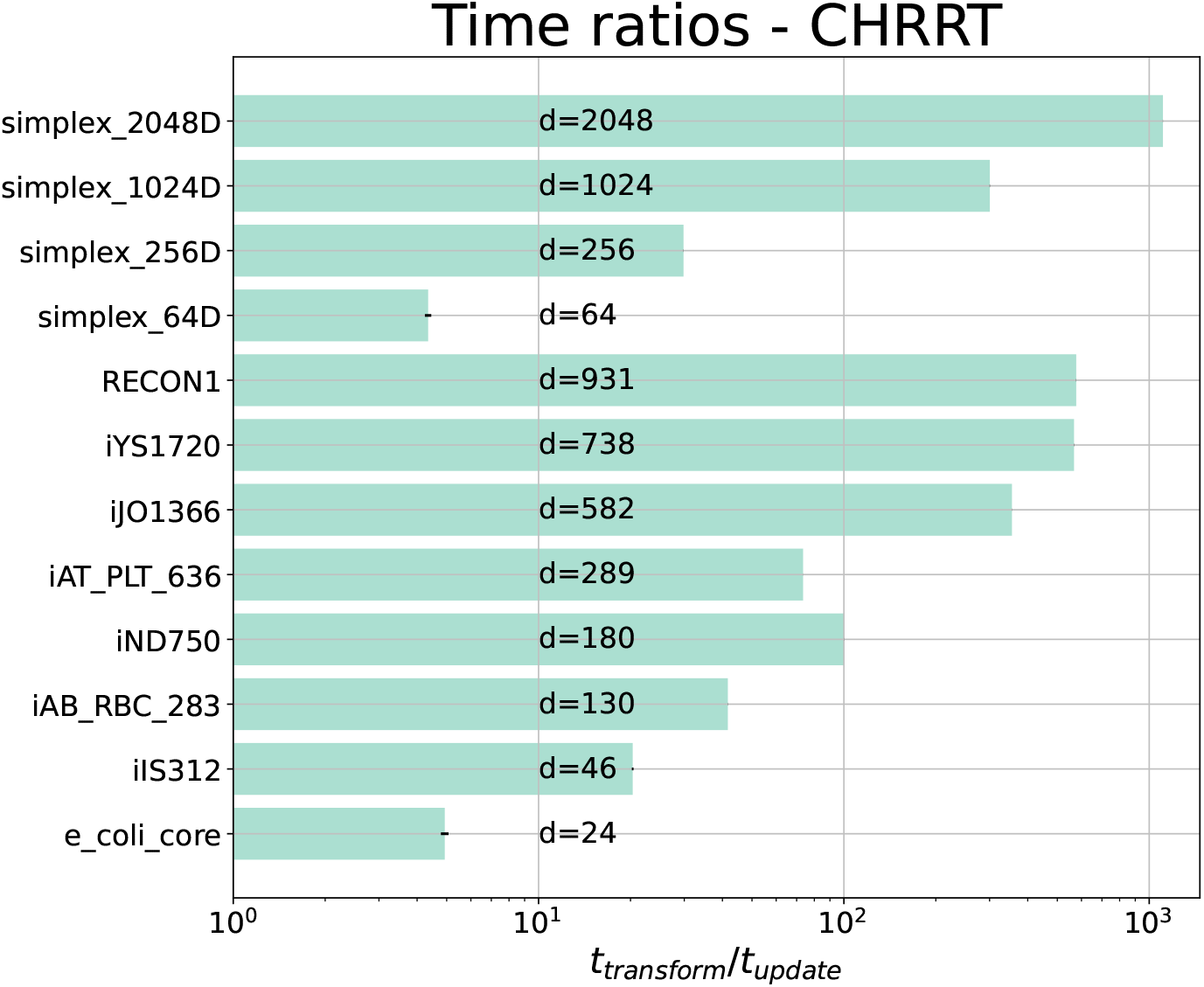
Mean time ratios of one back-transform *t*_transform_ vs. one CHRRT update *t*_update_ for simplices and GEMs (horizontal axis scaled logarithmically). Ratios indicate that in all cases, the cost to transform a sample back to the original flux space exceeds the cost of generating a sample by far. In all but the smallest test problems, the error bars of four runs are too small (< 1%) to be visible. For the simplices the ratio increases quadratically with dimension (0.00024 · *d*^2^ + 0.044 · *d* + 1.4). For GEMs, there is a strong trend towards an increase in ratio with model dimension. Details on the models are found in SI Table S.1.

Across all test problems, *t*_transform_ is generally considerably larger than *t*_update_, with a time ratio of at least four for the smallest tested models, but which starkly increases with the dimension of the sampling problem. Given such large time ratios, it is plausible to expect thinning to be beneficial for the sampling efficiency of CHRRT in the context of UCPS. However, only by considering the relation between the time ratios and the autocorrelation mediated by the *ESS*, which is specific to the combination of model characteristic and CHRRT, we are able to find out if thinning actually is practically beneficial. In conclusion, low autocorrelation should not be the only deciding factor when suggesting rules of thumb for thinning constants.

### 3.2 Benchmarking CHRRT from the perspective of thinning

To study how the performance of CHRRT depends on thinning for the two classes of test problems, we benchmarked the sampling efficiency, in terms of *ESS/t*, for a wide variety of thinning constants.

To find thinning constants that yield high *ESS/t* values, pre-runs with *τ* = *d* were performed for each test problem. From the pre-runs, we estimated appropriate ranges of problem-specific thinning constants and selected a set of at least five *τ* ‘s distributed over that range.

For simplices and GEMs, Fig. 2 shows the obtained *ESS/t*, relative to the unthinned CHRRT baseline. The plots fan out in a series of curves for the test problems with different model dimensions. As a rule, for higher dimensions, higher relative *ESS/t* values are achieved. All curves share the characteristic that with increasing *τ*, the relative *ESS/t* first increases before dropping again. The peak indicates the optimal thinning constant choice, specific to the investigated test problem. While for simplices pronounced peaks emerge, the curves for GEMs show plateaus, indicating that a range of thinning constants perform equally well. This apparent insensitivity in *ESS/t* performance for GEMs is convenient, since it implies that hyperparameter tuning of *τ* does not need to be precise for achieving near optimal *ESS/t* values, if its value is set to a suitable order of magnitude. Clearly, thinning constants at the lower end of the plateau are to be preferred due to a better resource usage (see below).

**Figure 2:**
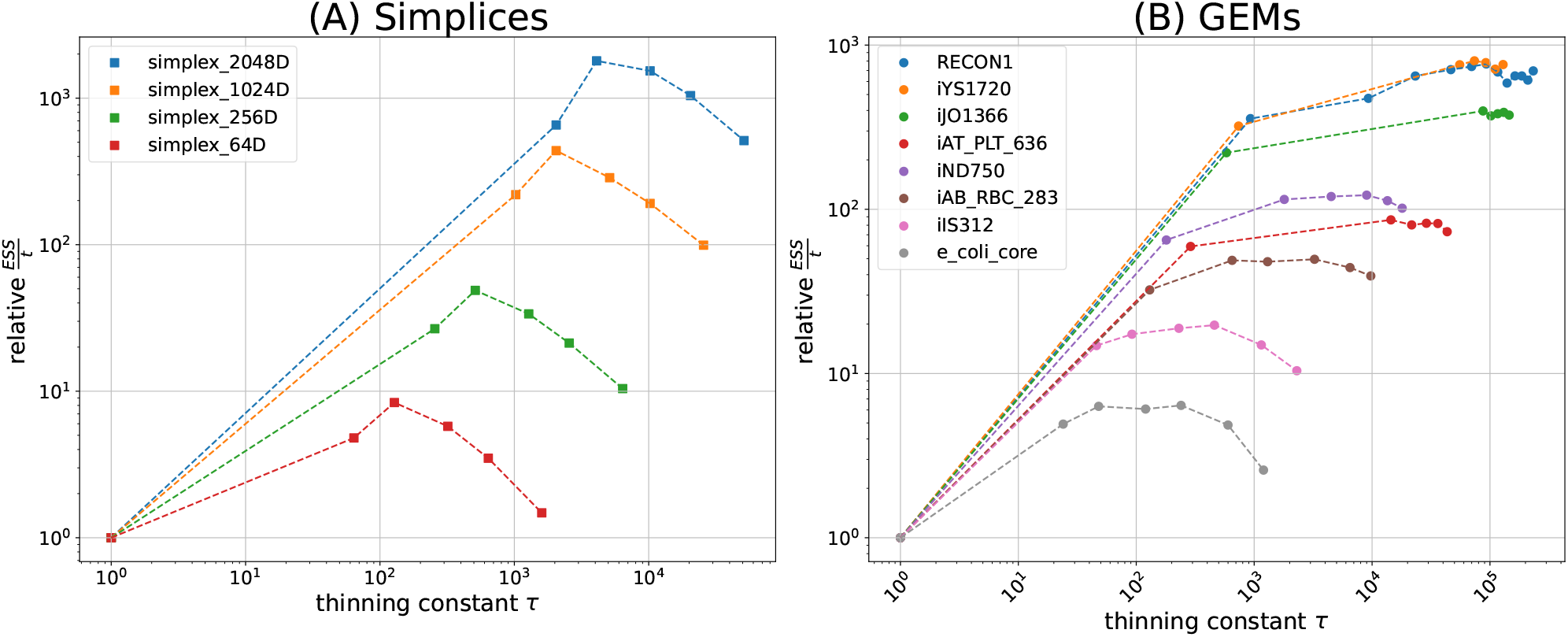
Double logarithmic plot of sampling efficiencies, relative to the unthinned sampling efficiencies, for selected thinning constants *τ*. The corresponding absolute *ESS/t* values are provided in SI Fig. S.1. (**A**) Benchmark results for simplices. The peak efficiency is reached for a thinning constant 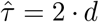, with *d* being the number of effective dimensions. Over all, the maximum speed-up is achieved for the 2,064-dimensional simplex, being 1,798 times faster in terms of *ESS/t* compared to unthinned CHRRT. (**B**) Benchmark results for GEMs. Setting *τ* > 1 generally improves the *ESS/t*. For increasing *d*, larger thinning constants further boost the sampling efficiency. In all cases, the emerging plateaus indicate that the optimal *ESS/t* is not overly sensitive to the precise choice of the thinning constant.

To study how the optimal (in terms of sampling efficiency) thinning constant relates to the problem dimension, we then quantified the speed-up of CHRRT for the apparent best thinning constant, hitherto denoted 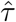, among the set of tested constants. Results are shown in Fig. 3(A). For simplices, relative speed-ups grow roughly to the square of the polytope dimension. The relative speed-ups achieved for GEMs also grow with effective model dimension, with values ranging from 6 for the *E. coli* core model up to 770 for *Recon1*. While CHHRT with 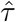 for *Recon1* required 0.32 s to converge and reach an *ESS* of 4,954, the unthinned variant required almost 80 times longer (25 s) to converge, while it reached an *ESS* of only 507. Notably, the speed-ups observed for GEMs surpass those observed for simplices of similar dimensions. Hence, our benchmarks show that the beneficial effect of thinning, when appropriately tuned, is not only plausible in theory, but it is also of high practical relevance for solving high-dimensional UCPS problems.

**Figure 3:**
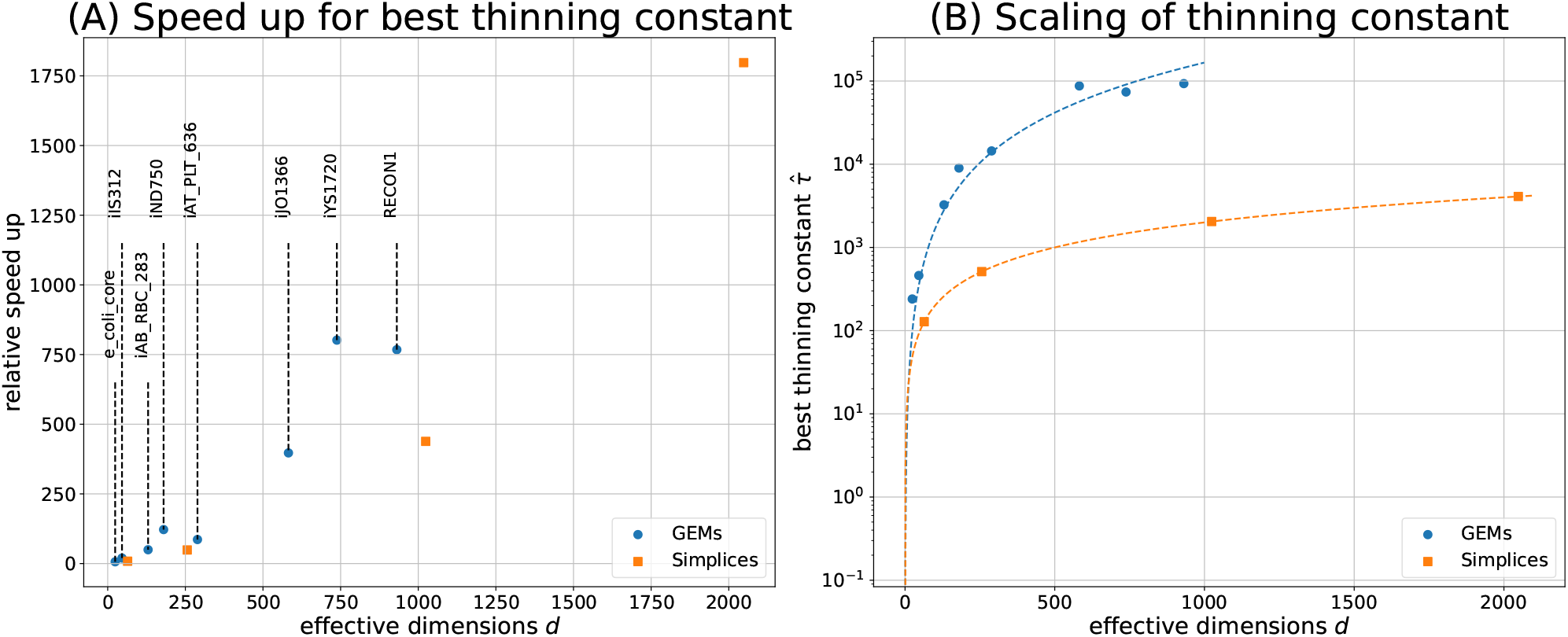
(**A**) Speed-ups achieved for the best thinning constants 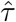, compared to the unthinned baseline, for simplices (orange squares) and GEMs (blue cicles). (**B**) Scaling of the best thinning constant given the effective model dimensions *d* (vertical axis scaled logarithmically). Guidance for the selection of useful thinning constants 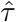 for GEMs (blue line) and simplices (orange line) is obtained by regression. For problem dimensions above 500, 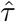 is about two orders of magnitude larger for GEMs than for simplices.

### 3.3 Simple thinning rules for simplices and GEMs

For *d*-dimensional simplices, the linear relation

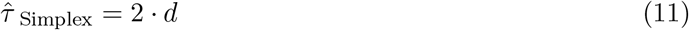

explains the measured data perfectly. For GEMs, on the other hand, a quadratic relation of the form 0.164 *· d*^2^ matches the data well. Considering Fig. 2(B), indicating that optimal sampling performance is not overly sensitive to thinning constant choice, as a memorable rule of thumb for efficient GEM sampling, we propose:

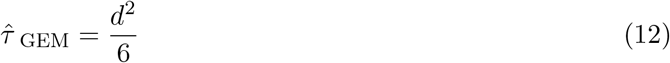

which only marginally deviates (less than 1.5%) from the fitted line.

Compared to the advice previously given by Haraldsdóttir et al. (2017), i.e., 8 · *d*^2^, our GEM guideline advocates thinning constants that are a factor of 48 smaller. To put the difference between the former and our updated guideline into perspective, a 48-fold increase in *τ* means a 48-fold increase in CHRRT update steps, of which the vast majority is discarded. As we have argued before, discarding samples can lead to an improvement of sampling efficiency, but only if the loss of information due to dropping samples is over-compensated by information from additional CHRRT update steps. In addition, in Fig. 2 we show that selecting the thinning constant too large risks missing the performance peak, which then wastes computational resources. In particular, we observe that a 48-fold increase in *τ* leads to lower sampling efficiencies for eight out of the twelve test problems. For the remaining four test-problems, i.e., GEMs of higher dimensionality, we did not attempt to characterize the plateau around the best thinning constants, since this would be of little practical relevance. It is generally preferable to set the thinning constant to the minimal value on the plateau, because then less work is discarded, and more samples are stored than for larger thinning constants, while achieving the same *ESS/t*. Nonetheless, it is imaginable that the performance plateaus for the four largest GEMs stretch across an even wider range of thinning constants, meaning that the difference in *ESS/t* resulting from the previous rule of Haraldsdóttir et al. (2017) and our guideline may turn out to be smaller.

While more work could be invested into fine-tuning 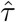 for the problem at hand, this risks investing more computational work than is eventually gained, because the estimation of *ESS* is noisy for short CHRRT runs. In summary, we showed that a previous guideline for GEM sampling is suboptimal from the perspective of sampling efficiency. Instead, we promote a new GEM guideline, given by Eq. (12), which empirically maximizes sampling performance without unnecessarily discarding computational work (green computing).

### 3.4 Guideline verification with the *Recon3D* GEM

To verify our proposed guideline for CHRRT sampling of GEMs, we applied it to an out-of-sample GEM. For this we selected *Recon3D* the largest GEMs currently available in the BiGG database, with an efficient dimension more than five times larger than the largest model for which benchmarks were performed (Supplementary Table S.1). Because application of CHRRT with *τ* = 1 is impractical for *Recon3D*, sampling efficiencies were compared using the previous and our newly proposed advice for thinning.

We benchmarked the previous advice of Haraldsdóttir et al. (2017) and observed that CHRRT requires 1.1 h to produce and store one sample per chain with *τ* = 8 · 4, 861^2^ = 23, 629, 321. Assuming optimistically that each of these samples is uncorrelated, i.e., *ESS* = 1, we project that it takes approximately 110 h until convergence, when four parallel chains are used. This corresponds, in the best of all cases, to an *ESS/t* of at most 0.001 s^−1^. Using the newly proposed guideline in Eq. (12), CHRRT with a thinning factor of ≈ 3, 938, 220, it takes 27 h to generate 1, 000 samples per chain.

Using four parallel chains, this setup results in an *ESS* of 726 with an 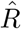. Therewith, our guideline yielded an *ESS/t* = 0.0075 s^−1^, which is 7.5 times more efficient than the overly optimistic upper bound we found for the previously advised rule. In summary, the new guideline is not only more efficient, but it also produces a converged set of samples for storage in a fraction of the time.

## 4 Conclusion and outlook

CHRRT is the leading algorithm for sampling convex polytopes uniformly, having a single adjustable parameter, the thinning constant. In this work, we showed that the choice of the thinning constant has dramatic consequences on sampling efficiencies, which are instrumental for solving high-dimensional UCPS problems. Here, selecting an appropriate thinning constant can make the difference between sampling success and failure, thus, being the key to sample challenging network models to convergence. Especially for contemporary GEM sampling, it is therefore crucial to have a guideline at hand for the selection of the thinning constant that achieves near optimal performance.

With a range of UCPS problems at hand, we studied the effect of thinning on CHRRT sampling performance quantitatively, after explaining the statistical underpinnings of the efficiency metric (*ESS/t*). From our quantitative benchmarks, we derived simple rules for optimal CHRRT thinning constant choice for simplices and GEMs. Applying these rules is not only beneficial for obtaining uniform samples of polytopes faster, but also helps to migrate UCPS towards green computing, as both the CPU time and the storage cost of samples are reduced. Concerning the question of how the computational requirements of CHRRT scale with effective model dimension, our numerical results reveal a quadratic and linear correlation for simplices and GEMs, respectively. Benchmarking new UPCS algorithms and comparing their performances with those of leading algorithms, such as CHRRT, is important to advance the field and a topic of active research (Chalkis et al., 2021b). Recognizing the substantial impact of thinning on the performance of CHRRT, we advise performing comparisons with tuned thinning, and to report the used thinning constant, which will help their reproduction.

Replacing the sequential per-sample by a one-shot or batched back-transform (algorithm 1, L 12) could be more efficient, as long as memory issues are not limiting (storing unthinned samples of large models, such as *Recon3D*, consumes prohibitively much memory). To combat memory bottlenecks, Stein thinning (Riabiz et al., 2022), a technique to optimally compress the MCMC output, may be applied before sample back-transformation. However, while this combination of techniques is promising, the implementation is not straightforward. In contrast, fixed-frequency thinning, as discussed in this work, is already implemented in many CHRRT packages and immediately boosts the performance of CHRRT for sampling polytopes uniformly.

Despite thinning being a common MCMC practice, it is still controversially discussed. Statisticians have pointed out that thinning is often not necessary and that it typically wastes computational resources, unless the cost of using the samples is high (Geyer, 1992; Link and Eaton, 2012). Even then, Geyer argues that a thinning constant of two or three times the problem dimension should be suited in nearly all cases. Consequently, the conventional advice given to the MCMC practitioner is to not thin MCMC outputs, unless memory or post-processing capabilities are practically limiting. By showing that CHRRT stands out among MCMC algorithms in that thinning is critical to performance for sampling high-dimensional convex polytopes, such as GEMs, our study encourages researchers to systematically examine conventional MCMC advice in their specific application domain.

## Supporting information

Supporting information on models, additional data and references

## Acknowledgements

The authors thank Axel Theorell, Richard D. Paul and Johannes Seiffarth for fruitful discussions and feedback.

