## Supporting information on models, additional data and references for "CHRRT: boosting coordinate hit-and-run with rounding by thinning"

### S.1 Benchmark Models

Table S.1: Overview of models and simplices used. The number of constraints gives the number of non-redundant constraints after preprocessing with PolyRound, i.e. after elimination of redundancies and 0-facets.

| Model | Type | #reactions | # effective polytope dimensions $d$ | #constraints $m$ | Reference |
|---|---|---|---|---|---|
| e_coli_core | GEM | 95 | 24 | 36 | Orth et al. (2010) |
| iIS312 | GEM | 519 | 46 | 137 | Shiratsubaki et al. (2020) |
| iAB_RBC_283 | GEM | 469 | 130 | 188 | Bordbar et al. (2011) |
| iND750 | GEM | 1,266 | 180 | 273 | Duarte et al. (2004) |
| iAT_PLT_636 | GEM | 1,008 | 289 | 593 | Thomas et al. (2014) |
| iJO1366 | GEM | 2,583 | 582 | 752 | Orth et al. (2011) |
| iYS1720 | GEM | 3,357 | 738 | 962 | Seif et al. (2018) |
| RECON1 | GEM | 3,741 | 931 | 2,120 | Duarte et al. (2007) |
| Recon3D | GEM | 10,600 | 4,861 | 11,195 | Brunk et al. (2018) |
| simplex_64D | Simplex | | 64 | 65 | |
| simplex_256D | Simplex | | 256 | 257 | |
| simplex_1024D | Simplex | | 1,024 | 1,025 | |
| simplex_2048D | Simplex | | 2,048 | 2,049 | |

### S.2 Measured $ESS/t$

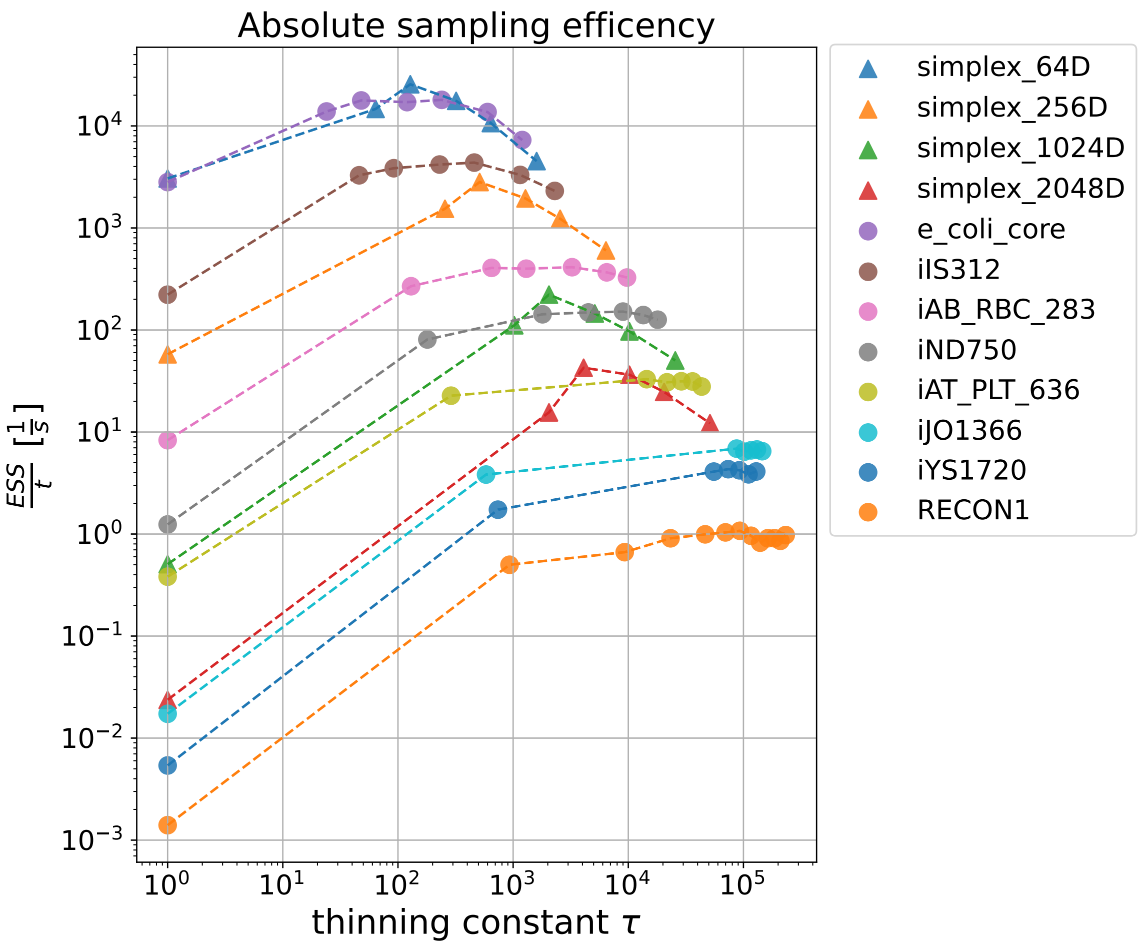


Figure S.1: Double logarithmic plot of absolute sampling efficiencies as measured by $ESS/t$ for selected thinning constants $\tau$. Note that CHRRT for simplices achieves a higher absolute performance compared to sampling GEMs with comparable dimensions. As an example, the best thinning constant $\hat{\tau}$ for simplex_2048D lies at around the same value as the peak for the iAT_PLT_636 model, which has 582 dimensions. By comparing the absolute $ESS/t$ across various problems, it becomes clear that the effective dimensionality of the polytope is one, but not the only component determining sampling complexity.
